# When an insecticide affects the adaptive value of intraguild predation by an invader

**DOI:** 10.1101/349852

**Authors:** Paula Cabrera, Daniel Cormier, Marianne Bessette, Vanessa Cruz, Éric Lucas

**Author notes:** Corresponding author (PC). These authors contributed equally to this work (Paula Cabrera, Daniel Cormier, Éric Lucas). These authors also contributed equally to this work (Marianne Bessette, Vanessa Cruz).

## Abstract

Biological invasions can generate major ecological disturbances, such as changes in species diversity and structure of communities. It is believed that the multicolored Asian ladybeetle, *Harmonia axyridis* Pallas (Col, Coccinellidae), recognized as one of the most invasive insects in the world, has reduced native coccinellids populations in several areas and is considered as a threat for biodiversity at large. A significant trait, favoring its invasiveness and its dominance over indigenous ladybeetles, is intraguild predation (IGP). IGP has advantageous adaptive value for individuals, removing competitors, potential predators and providing an alternative nutritive resource, when main resources are scarce. Previous research demonstrated that this invasive ladybeetle is highly susceptible to the reduced-risk insecticide novaluron, a chitin synthesis inhibitor, whereas the North American indigenous competitor, *Coleomegilla maculata* DeGeer (Col, Coccinellidae), is not. Our study explores the adaptive value of IGP for each of the two coccinellids after preying on each other’s larvae, previously treated with insecticide. Our first hypothesis is that the invasive ladybeetle, susceptible to the insecticide, should lose the adaptive value of IGP, while the native predator not. Our second hypothesis is that the adaptive value of IGP for the invasive predator will be recovered over time, as a result of neutralisation of the insecticide by the intraguild prey (native species). The results support both hypotheses, and show that an insecticide can completely remove the adaptive value of IGP for the invader, while it does not change for the indigenous ladybeetle. Moreover, the study demonstrates that if the intraguild prey (non-susceptible to the insecticide) undergoes molt after being exposed to the insecticide, the adaptive value for the intraguild predator is restored.

## Introduction

Biological invasions can interfere with several ecological and evolutionary processes [1], such as changes in species diversity and structure of communities [2], affecting ecosystem services and causing huge economic losses [3–5]. Some of the traits that characterize invasive species are high adaptability to changes in the environment, high fecundity and dispersion[6, 7], and broad diet range [8–10]. Another factor contributing to the success of invaders is competitive ability, and more specifically intraguild predation (IGP) [11–14]. IGP, is defined as the predation on a competitor [15], and confers adaptive advantages for of the intraguild predator. These include removal of competitors, removal of potential predators, and an alternative nutritive resource when main resources (extraguild prey) are scarce. Therefore, IGP has important consequences for the distribution, abundance, and evolution of species involved (intraguild predator, intraguild prey and shared resource) [16] as well as for applied topics such as conservation management and biological control [17, 18].

The multicolored Asian ladybeetle, *Harmonia axyridis* Pallas (Col., Coccinellidae), widely recognized as one of the most invasive insects on the world [10, 19, 20], has been introduced in numerous ecosystems worldwide [19, 21–29]. As an invasive species, it has caused adverse impacts on the wine industry [30–32] and as a household invader during winter. It is an effective generalist predator of numerous hemipteran pests [33], with higher fertility and fecundity than competitive ladybeetle species [27, 34–35], significant adaptability and resilience [36], and has a dominant role as an intraguild predator [37–41]. These attributes suit the multicolored Asian ladybeetle to be a top predator according to several experts [42]. Consequently, it has reduced diversity of natural enemies in certain regions, especially ladybeetles, [33] through exploitative competition and intraguild predation [34, 39, 43–44].

The twelve spotted ladybeetle, *Coleomegilla maculata* (DeGeer) (Col., Coccinellidae), is one of the sympatric competitors of *H. axyridis*, in Nearctic apple orchards [45] as well as in other agricultural ecosystems, such as potato [46–47] and sweet corn [24]. This ladybeetle, is an indigenous generalist predator in North America, and an important biocontrol agent in various crops [45, 47–49] that also participates in IGP with other coccinellids. Thus, both coccinellids, *C. maculata* and *H. axyridis*, frequently engage in mutual IGP [50].

Although IGP has an important adaptive value for the intraguild predator, it also involves certain risks, such as the risk of injuries or an increase of exposure to pesticides [51]. Moreover, it has been proposed that 75 % of the biodiversity can be concentrated in agricultural landscapes and thus, species are exposed to pesticides used for plant protection [52]. Despite of this, very few studies have addressed the impact of pesticides on IGP [51, 53]. Besides, it remains difficult to establish how pesticide applications may affect success of invasive species using IGP as competitive arm.

Previous research has shown that novaluron (Rimon^®^ EC 10), a reduced-risk insecticide (RRI) and chitin synthesis inhibitor [54], used to control the codling moth, *Cydia pomonella* (L.) (Lep., Tortricidae), a major pest in Quebec apple orchards [55], has differential lethal and sublethal effects on *H. axyridis* compared to *C. maculata*, the multicolored Asian ladybeetle being drastically more susceptible to the insecticide than its indigenous intraguild competitor [56–57]. Insects can overcome the effects of toxic compounds by tolerating them or neutralizing them by several mechanisms, such as sequestration, excretion and detoxification [58], and these can determine differences in susceptibilities to pesticides among natural enemies [59].

The aims of this study are 1) to compare the adaptive value of mutual IGP, for the two coccinellids, the invasive and the native species, when the IGP prey faced exposition to an insecticide used in the invaded area; and 2) to investigate if the effect of the exposition on the adaptive value of IGP can disappear over time through neutralisation of the insecticide by the intraguild prey, the least susceptible species (the indigenous) to novaluron. We hypothesize that the adaptive value of IGP, for the indigenous tolerant twelve spotted ladybeetle, will be maintained in the presence of the insecticide, whereas it will be greatly reduced for the susceptible invasive Asian ladybeetle. We also hypothesize that the adaptive value of IGP for the susceptible invasive species will be recovered, after a period, when the native IG prey will have neutralized the insecticide.

## Materials and methods

### Insects

Laboratory rearings of *C. maculata* and *H. axyridis* originated from ladybeetle adults collected in the field in Quebec in 2016 at Sainte-Agathe (46°23′0.3′′N and 71°24′33.5′′W). New individuals coming from the field were added every year to the rearings. These were in a growth chamber at 24°C, 16L: 8D, and 70% RH at Laboratoire de Lutte Biologique from the Université du Québec à Montréal. Insects were provided with pollen, sweetened water solution (10 % sugar), and green peach aphids, *Myzus persicae* (Sulz.) (Hem., Aphididae), reared on potato plants, *Solanum tuberosum* L.

### Insecticide

Rimon^®^ 10 EC (Makhteshim Agan of North America, Raleigh, NC) field rate recommended for apple orchards in Quebec (100 g a.i. ha^−1^) and a spray volume of 1000 litres ha^−1^, most used by apple growers in Quebec, were considered in the concentration for bioassays: 100 mg a.i. litre^−1^ novaluron.

### Bioassays

A first bioassay was intended to assess the effect of novaluron on intraguild predation by both coccinellid species on the other one. A second bioassay was performed to explore neutralisation of pesticide by *C. maculata* as an intraguild prey, and its effect on *H. axyridis* as the intraguild predator.

### Effect of the insecticide on intraguild predation

Experimental units consisted in a newly molted second instar *C. maculata* or *H. axyridis* larvae (intraguild predator), individually weighted, and placed in 5 cm Petri dishes with 5 first instar larvae from the opposite ladybeetle species (intraguild prey), previously treated with novaluron or water with a Potter Tower (1 ml solution and 2 mg cm-2 aqueous insecticide deposit) and euthanized by freezing to avoid cannibalism. Also thawed for 3 min at 20°C before the bioassay) (Fig 1). Each experimental unit was replicated 17 times. Intraguild predators and prey were left together during 24 h. Leftovers of intraguild prey were noted and removed afterwards. After that, intraguild predators were fed with *M. persicae* aphids daily until the adult stage. Mortality and stage of development were noted on a daily basis. Once the individuals became adults, they were weighted again.

**Figure 1.**
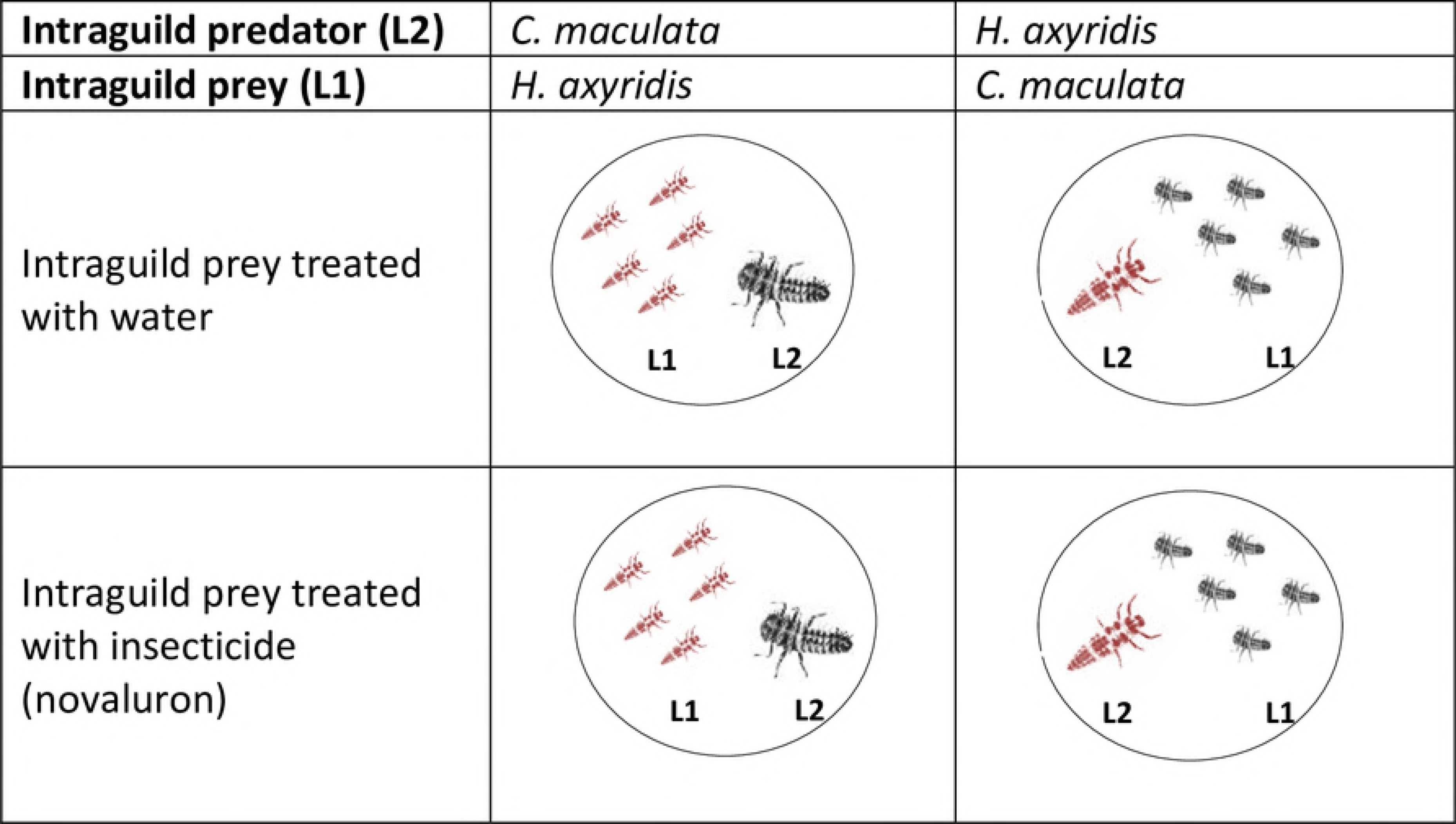
Experimental design used to assess the effect of novaluron on IGP. Each experimental unit consisted of a second instar larva (intraguild predator) placed with five 1st instar newly hatched larvae from the opposite coccinellid species. Each experimental unit was replicated 17 times.

### Neutralisation of the insecticide by the indigenous intraguild prey, *C. maculata* and its effect on the invasive intraguild predator *H. axyridis*

Three treatments, representing three situations of neutralisation of the insecticide in the intraguild prey, were considered in this bioassay:

a. **T0:** 1^st^ instar larvae of the indigenous ladybeetle or aphids, insecticide or water treated, and immediately euthanized,
b. **T-24h:** first instar larvae of the indigenous ladybeetle or aphids, insecticide or water treated and euthanized 24 h after,
c. **T-Molt:** first instar larvae of the indigenous ladybeetle or aphids, insecticide or water treated and euthanized after molt to 2^nd^ instar.

Additionally, three treatments were associated to each of the treatments mentioned above;

1. **XG-Water**: Aphids treated with water, representing optimal extraguild prey, and offered *ad libitum* to intraguild predators,
2. **IG-Water**: 1^st^ instar larvae of the indigenous LB, treated with water, and
3. **IG-Insecticide**: 1^st^ instar larvae of the indigenous LB, treated with the insecticide (novaluron).

A Potter tower was used to treat larvae (1 ml solution and 2 mg/cm^2^ aqueous insecticide deposit). All preys were weighted prior to freezing. Experimental units were replicated 17 times and consisted of five *C. maculata* larvae, thawed during 3 min at 20°C, and placed in a 5 cm Petri dish with a newly molted 3^rd^ instar *H. axyridis* larva, previously starved during 24 h. Third instar larvae were used as intraguild predators in this bioassay, in order to encourage IGP by reducing the prey/predator body mass ratio. Intraguild predators were in contact with extraguild and intraguild prey during 24 h. Intraguild and extraguild prey leftovers were noted and removed afterwards and intraguild predators were fed with fresh green peach aphids daily until reaching the adult stage. Mortality was assessed daily and *H. axyridis* individuals were weighted after reaching adult stage.

### Calculations and statistical analyses

Differences in frequencies of mortality in both bioassays were evaluated with a Chi^2^ test (α = 0.05) [60], and pairwise comparisons, with Bonferroni corrections of the p value, were performed to detect differences among treatments [61]. Results are presented as mortality percentages. Survival was analysed by means of Survival Analysis and a Proportional Hazard model to detect differences among groups [62].

Voracity of the intraguild predator was calculated by assessing 1^st^ instar larvae (intraguild prey) leftovers after 24 h of IGP. Mass of consumed prey (mg) was estimated from percentages of leftover prey and weight of prey before freezing (for the neutralisation of the insecticide by the intraguild prey section). Weight increase of intraguild predators until the adult stage was calculated subtracting second or third instar larvae initial weight from adult weight (g). This variable, as well as voracity, were analysed with the independent samples Student’s t test (α = 0.05) to investigate the effect of novaluron on both ladybeetle species and with Analysis of Variance and the Tukey test or the non-parametric Wilcoxon test and the Steel-Dwass All Pairs test (α = 0.05) (SAS Institute, 2015) to detect differences among treatments in the case of the effect of insecticide neutralisation by intraguild prey on *H. axyridis*, since data did not always meet the normality assumption. Time of development of *C. maculata* as the intraguild predator, in the first bioassay, was examined with the independent samples Student’s *t* test (α = 0.05) [59] and time of development of *H. axyridis* in the second bioassay was investigated with the Wilcoxon test (α = 0.05) [63]. JMP software v.12.1 (SAS Institute, Cary, NC) was used to perform all statistical analyses.

## Results

### Effect of the insecticide on intraguild predation

#### Voracity of intraguild predators

There were no differences in voracity between the insecticide treatment and the control, *t*(_30.99_) = 1.469, *p* = 0.152 for the indigenous ladybeetle. A different trend was found for *H. axyridis*. Less prey was ingested in the insecticide treatment than in the control *t*_(29.37)_ = -2.283, *p* = 0.0299 (Table 1).

**Table 1.** Voracity of intraguild predators after 24 h feeding on intraguild prey treated with water or insecticide. Comparisons within lines. Student t test (*p* < 0.05)

| | Intraguild prey consumed (% ± SE) | |
| --- | --- | --- |
|  | Control | Insecticide |
| Indigenous ladybeetle | 42.81 ± 7.27 A | 58.06 ± 7.41 A |
| Invasive ladybeetle | 61.56 ± 7.33 a | 40.00 ± 5.96 b |

#### Mortality-survival of intraguild predators

After feeding on intraguild prey (first instar invasive ladybeetle) treated with novaluron, mortality of indigenous ladybeetle second instar larvae, was not different than the control, *X*^2^_(1)_ = 0.013, *p* = 0.9086 (Fig 2-a). Surviving individuals molted normally and completed development until the adult stage. In the case of the invader *H. axyridis*, all larvae died after IGP in the novaluron treatment, which mortality was significantly higher than the control, *X*^2^_(1)_ = 15.245, *p* < 0.0001 (Fig 2-b). Survival analysis for intraguild predators, preying on intraguild prey treated with the insecticide, reveals a significant difference between survivals of both species over time, *X*^2^_(1)_ = 12.731, *p* = 0.0004. All *H. axyridis* larvae died between the first and fifth day following IGP and none of the predators were able to molt before death, whereas 70% of indigenous *C. maculata* individuals reached the adult stage (Fig 2-d).

**Figure 2.**
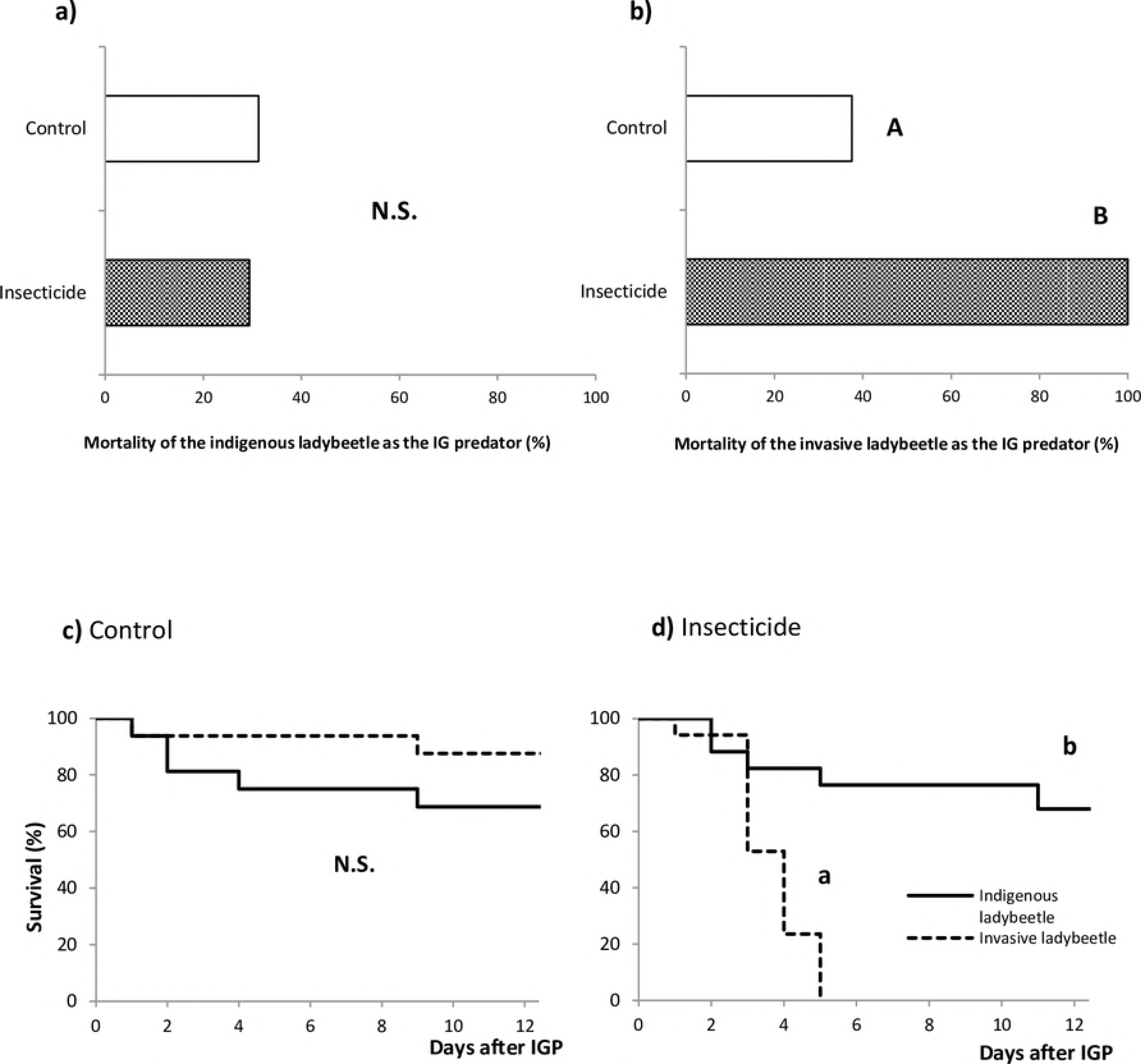
a) Mortality of the indigenous ladybeetle as the intraguild predator, b) Mortality of the invasive ladybeetle as the intraguild predator, after 24 h exposure of 2nd instar larvae (intraguild preys) to water or insecticide-treated preys. Chi^2^ (p < 0.05), c) Survival and time of death of intraguild predators after ingestion of intraguild prey treated with water and d) treated with the insecticide. Kaplan-Meier survival curves (p < 0.05).

#### Time of development and weight increase (indigenous ladybeetle)

Since all *H. axyridis* individuals in the insecticide treatment died at the second larval instar, results of time of development and weight increase were obtained only for *C. maculata* in this treatment. Time of development until adult stage, after IGP, for the twelve spotted ladybeetle, in the insecticide treatment, was not different than the control *t*_(21)_ = 0.622, *p* = 0.541 (Fig 3-a).

**Figure 3.**
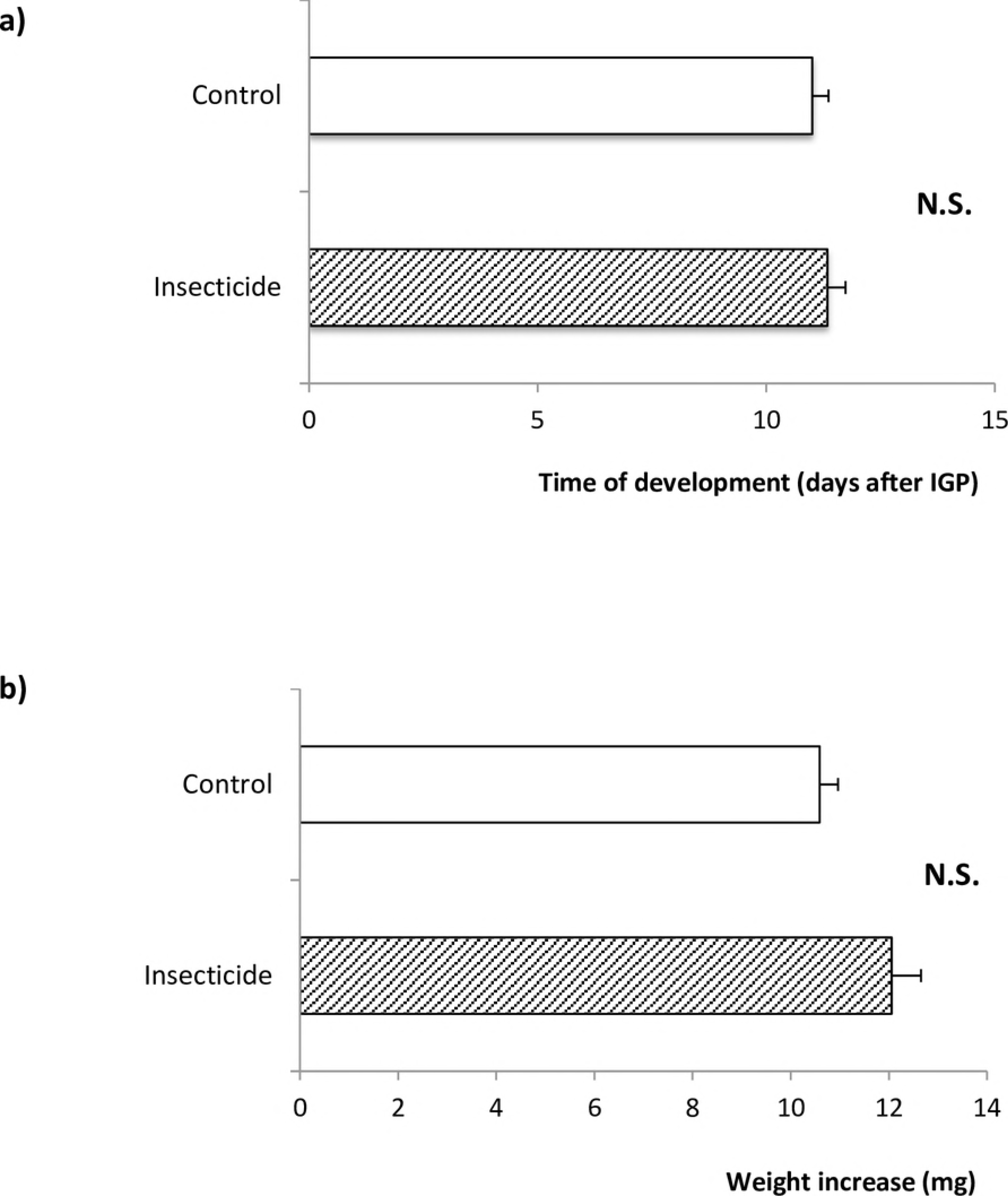
a) Time of development (days + SE) and b) weight increase (mg ± SE) of the indigenous ladybeetle, from 2nd instar larvae until adult stage, after 24 h feeding on intraguild prey treated with water or insecticide. Student t test (*p* < 0.05)

Weight increase was not statistically different between the control and novaluron treatments, *t*_(17.97)_ = 2.08, *p* = 0.0521 (Fig. 3-b).

### Neutralisation of the insecticide by the indigenous intraguild prey and its effect on the invasive intraguild predator

#### Voracity of the invasive ladybeetle

*Harmonia axyridis* consumed similar proportions of intraguild prey treated with water in the three treatments: *X*^*2*^_(1)_ = 5.004, *p* = 0.082. When comparing voracity of *H. axyridis* preying upon *C. maculata*, treated with novaluron, among the three treatments, there were slightly more intraguild preys consumed in the T0 treatment compared to the other treatments. However the T-24h treatment was not different from T-Molt: *F*_(2)_ = 11.12, *p* = 0.0002 (Table 2.).

**Table 2.** Voracity of *H. axyridis* after 24h preying upon extraguild (XG) and intraguild (IG) prey treated with insecticide or water. Comparisons are done within lines. ANOVA or Wilcoxon test (*p* < 0.05).

| Treatment | Consumed prey (mg ± SE) |  |  |
| --- | --- | --- | --- |
|  | T0 | T-24h | T-Molt |
| XG-Water | 7.37 ± 1.95 **a** | 9.76 ± 1.20 **a** | 7.79 ± 0.93 **a** |
| IG-Water | 0.77 ± 0.10 **A** | 0.92 ± 0.17 **A** | 1.16 ± 0.18 **A** |
| IG-Insecticide | 1.07 ± 0.05 **A'** | 0.40 ± 0.04 **B'** | 0.69 ± 0.16 **B'** |

### Mortality-survival of the invasive ladybeetle as the intraguild predator

Mortality of *H. axyridis* was not different among the three neutralisation treatments (T0, T-24h and T-Molt), when consuming water-treated extraguild prey, this is green peach aphids: *X*^*2*^_*(2)*_ = 0.31, *p* = 0.851. Similarly, no differences were found among treatments when intraguild predators consumed *C. maculata* first instar larvae treated with water, *X*^*2*^_*(2)*_ = 0.857, *p* = 0.652. However, when comparing mortality frequencies of the invasive ladybeetle after feeding on the indigenous ladybeetle treated with novaluron, the T-Molt treatment showed significantly less mortality *X*^*2*^_*(1)*_ = 23.221, *p* < 0.0001; T-24h vs T-Molt: *X*^*2*^_*(1)*_ = 13.55, *p* = 0.0002; T0 vs T-Molt: *X*^*2*^_*(1)*_ = 13.38, *p* = 0.0003 (Bonferroni α = 0.025). Moreover, it was found that intraguild predators feeding on prey treated with novaluron in treatments T0 and T-24h had a higher mortality than their controls of extraguild prey and intraguild prey treated with water (*X*^*2*^_*(1)*_ = 20.285, *p* < 0.0001; T0-IG-Insecticide vs T0-XG-Water: *X*^*2*^_*(1)*_ = 14.4, *p* = 0.0001; T0-IG-Insecticide vs T0-IG-Water: *X*^*2*^_*(1)*_ = 17.14, *p* < 0.0001 and *X*^*2*^_*(1)*_ = 19.217, *p* < 0.0001; T-24h-IG-Insecticide vs T-24h-XG-Water: *X*^*2*^_*(1)*_ = 16.22, *p* < 0.0001; T-24h-IG-Insecticide vs T-24h-IG-Water: *X*^*2*^_*(1)*_ = 13.55, *p* = 0.0002, Bonferroni α = 0.017 respectively) (Fig 4-a).

**Figure 4.**
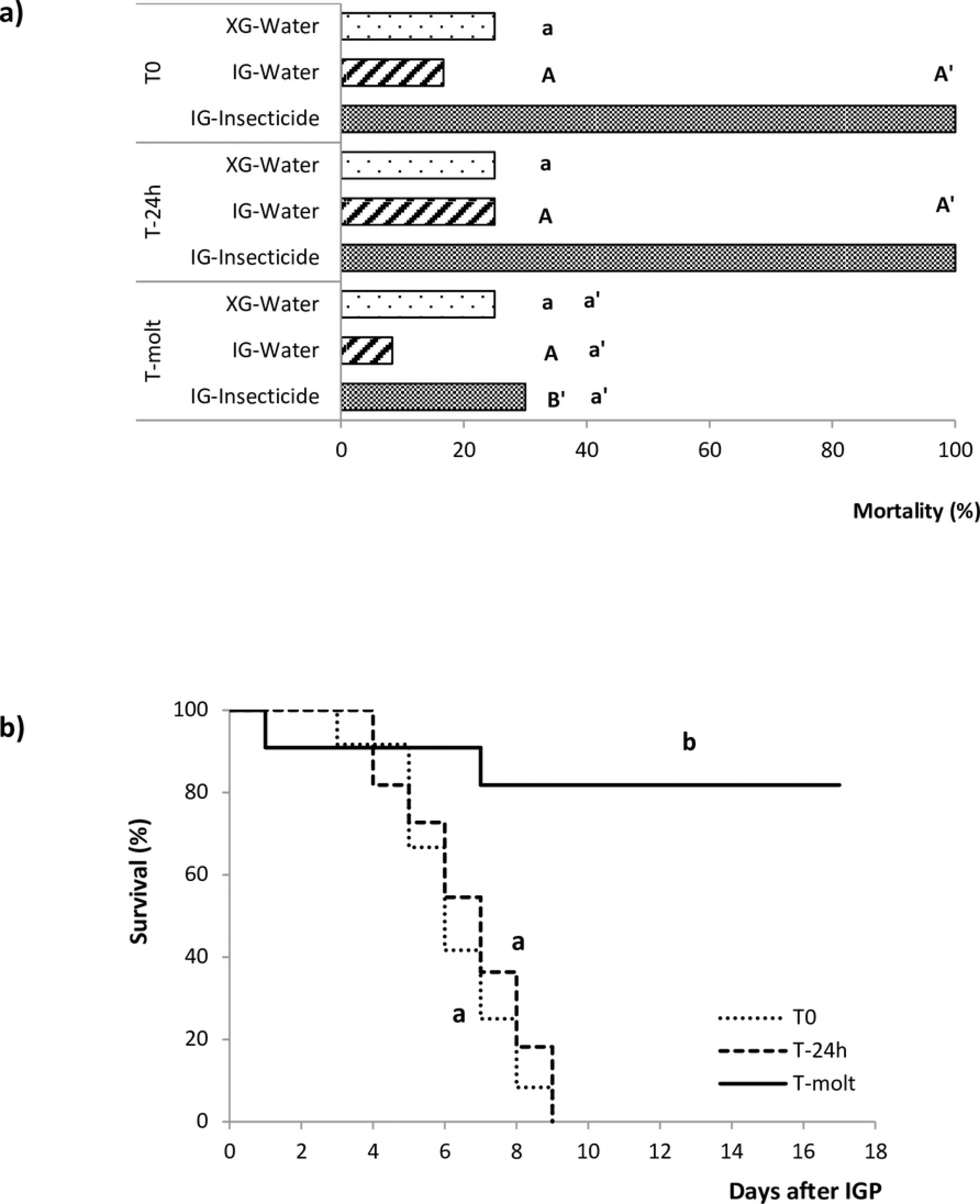
Experimental design used to assess the effect of novaluron on IGP. Each experimental unit consisted of a second instar larva (intraguild predator) placed with five 1st instar newly hatched larvae from the opposite coccinellid species. Each experimental unit was replicated 17 times.

A comparison of survival of the invasive ladybeetle over time among the three neutralisation treatments, after IGP on prey treated with novaluron, showed that 70 % of individuals in the T-Molt treatment became adults, whereas all individuals died in treatments T0 and T-24h before nine days (Fig 4-b). The Proportional Hazard Model indicated that this difference was significant: *X*^*2*^_*(2)*_= 19.91, *p* < 0.0001. Moreover, individuals in T0/IG-Insecticide were ° 15 times more prone to die than individuals in T-Molt/IG-Insecticide (*p* < 0.0001). Similarly, larvae in T-24h/IG-Insecticide had ° 11 times probabilities of death than larvae in T-Molt-IG-Insecticide (*p* = 0.0002).

### Time of development and weight increase of the invasive ladybeetle

Data from T0 and T-24h with treated intraguild preys are absent, since no individual reached the adult stage. No significant differences among treatments were found in the time of development after IGP, from third instar to adult stage: *X*^*2*^_(6)_ = 1.16, *p* = 0.98 (Fig 5-a).

**Figure 5.**
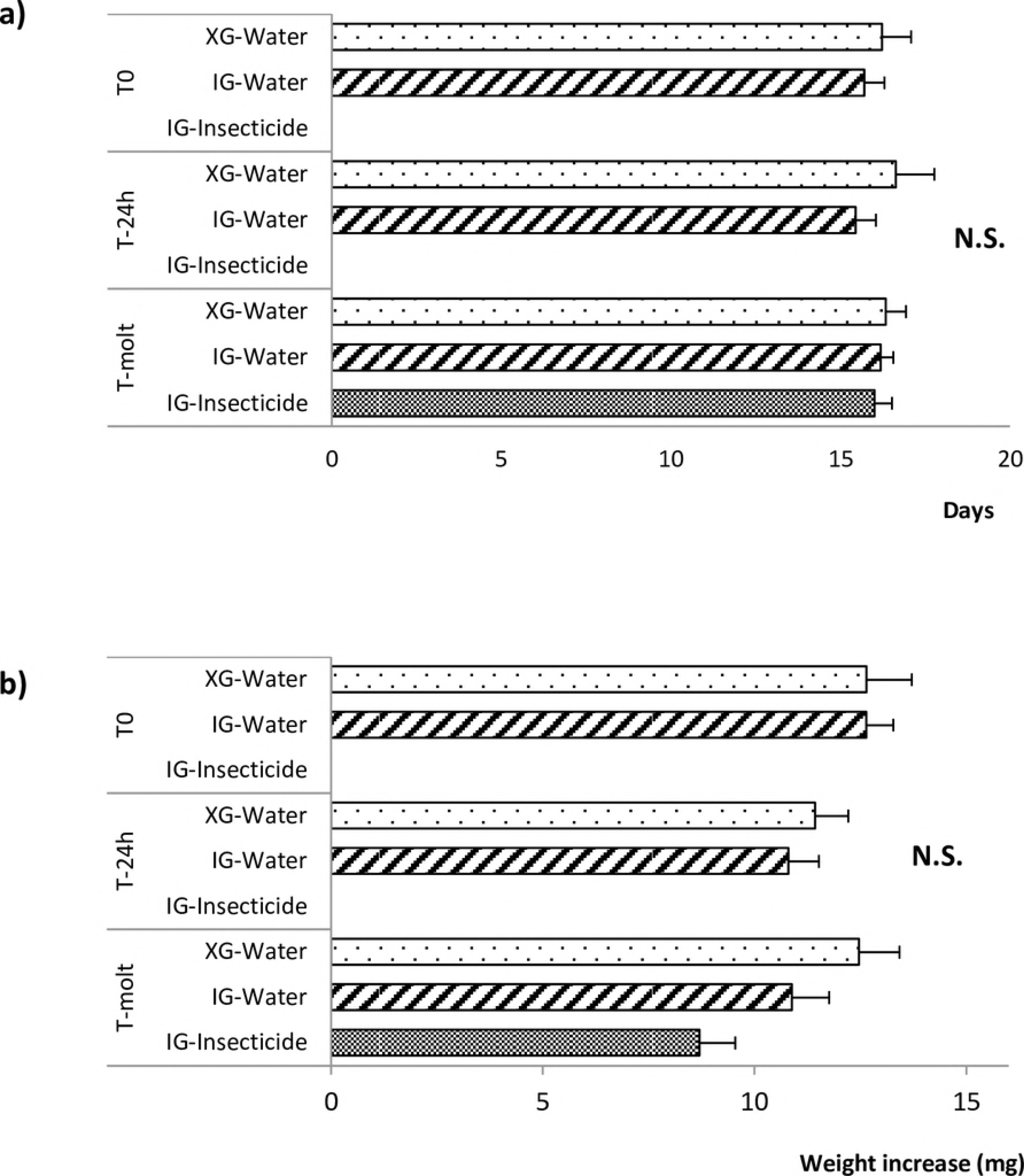
a) Time of development (days) of the invasive *H. axyridis*, from 3^rd^ instar larvae until adult stage, after 24 h feeding on intraguild prey treated with water or insecticide and extraguild prey; b) Weight increase (mg) of *H. axyridis* from third instar larvae until adult stage, after 24 h feeding on intraguild prey treated with insecticide or water and extraguild prey. Wilcoxon test (α = 0.05). **T0**: 1^st^ instar larvae of the indigenous ladybeetle or aphids, insecticide or water treated, and immediately euthanized; **T-24h:** first instar larvae of the indigenous ladybeetle or aphids, insecticide or water treated and euthanized 24 h after; **T-Molt:** first instar larvae of the indigenous ladybeetle or aphids, insecticide or water treated and euthanized after molt to 2^nd^ instar; **XG-Water:** Aphids treated with water; **IG-Water**: 1^st^ instar larvae of the indigenous ladybeetle, treated with water; **IG-Insecticide:** 1^st^ instar larvae of the indigenous ladybeetle, treated with insecticide (novaluron).

Similarly to time of development, no significant differences were detected in weight increase among treatments: *X*^*2*^_(6)_ = 11.93, *p* = 0.064 (Fig 5-b).

## Discussion

### Effect of the insecticide on the adaptive value of IGP for the invasive and the indigenous predators

Our results support our 1^st^ hypothesis that the adaptive value of IGP, for the invasive multicolored Asian ladybeetle, is completely lost when preying on intraguild prey treated with the insecticide. Furthermore, it is in accordance with previous research, confirming a differential susceptibility of both predator species to the reduced risk insecticide novaluron [56, 57]. At the opposite, after preying on intraguild insecticide-treated prey during 24 h, the adaptive value of IGP was maintained for the tolerant indigenous twelve spotted ladybeetle. Survival success was not different than survival in absence of the insecticide and neither voracity and weight increase, nor developmental time, were altered.

IGP is particularly advantageous for the adaptive value of intraguild predators, by removing competitors and potential predators, gaining energy and nutrition and consequently surviving, developing and reproducing when extraguild prey densities are low [16]. For instance, pollen is consumed by coccinellids at the beginning of the growing season in temperate regions, when prey densities are low, but in general it does not allow maturation of ovaries, therefore IGP is beneficial at that period [39]. Additionally, by the end of the season, IGP can help immature stages of predators to reach the adult stage, the overwintering stage in ladybeetles [64, 65]. However, our results show that an insecticide can drastically alter the outcome of IGP and even, completely cancel out its adaptive value for the intraguild predator.

Intraguild prey containing novaluron do not deter the invasive ladybeetle from consuming it and therefore, mortality is devastating for this insect after being exposed to contaminated prey for only 24 h. Which means that, in presence of novaluron, IGP is no longer an advantage for this predator. The adaptive value is completely lost and IGP becomes a highly risky behaviour, which it is not the case for the indigenous *C. maculata.* Thus, consequences of IGP between both, the indigenous and the invasive ladybeetles are reversed by the insecticide.

At a population level, a shift in the dominance of species on the coccinellid assemblage might be expected, particularly under conditions favoring IGP, such as scarcity of extraguild prey [66, 67] or habitats with little structure [68–69]. In this context, an insecticide might reduce populations of an invasive species, releasing populations of indigenous coccinellids as well as other natural enemies. Declines of natural enemies of Hemipterans, especially indigenous lady beetles in several regions where the multicolored Asian ladybeetle has established, have been attributed to interspecific competition for resources and the strong intraguild predation abilities of this invasive species [13]. Yet, the outcome of these repercussions in communities will depend on susceptibilities of intraguild members to the toxic compound, in populations.

### Neutralisation of intraguild prey and its effect on the intraguild predator

Our second hypothesis stating that the adaptive value of IGP, lost as a consequence of intraguild prey contamination by an insecticide, is recovered over time, is also supported by the results. This might be linked to neutralisation of the compound by the intraguild prey. Survival of the intraguild predator is significantly higher after feeding on contaminated intraguild prey having molted to the next stage compared to intraguild prey more recently treated. Results of voracity show that this finding is not due to a lower consumption of treated intraguild prey compared to the T-24h treatment. Weight increase of the intraguild predator, after predation on treated intraguild prey does not differ from weight increase after consumption of extraguild prey or intraguild prey treated with water in the highest intraguild prey neutralisation situation. The same trend is observed for the time of development of the intraguild predator. Therefore, benefits of intraguild predation are recovered over time.

Several phenomena may explain predator susceptibility to insecticides, among these, penetration, absorption and neutralisation may be significant. Penetration through the cuticle and rate of absorption of the compound in the arthropod body following topical contact or gut wall after ingestion may differ among species [54, 59]. Neutralisation can also be different among species. Toxic compounds can be neutralized by insects through sequestration (storage of compounds in an unaltered form), increased rates of excretion (removal of the toxic substance without altering its integrity) and detoxification (biochemical transformation of the compound in a way that it won’t harm the insect) [57]. Although we do not know how these mechanisms are involved, we formulate three hypotheses that might explain the recovery of the adaptive value of IGP by the intraguild predator over time: a) low penetration of the insecticide through the cuticle/gut wall of the intraguild prey, which is shed during the next molt or/and b) physiological neutralisation of the insecticide by the intraguild prey, occurring over time, c) elimination via feces. Thus, the outcome of IGP for the intraguild predator depends on the time that has passed, after exposure of the intraguild prey to the insecticide, specifically the time that is required for the intraguild prey to shed the contaminated cuticle or/and neutralise the insecticide to levels that will not harm the predator. The ability of *C. maculata* to neutralize the insecticide could be explained by a preadaptation to deal with toxic secondary plant compounds, since this species seems to be more phytophagous than *H. axyridis* [49, 70]. This hypothesis has already been mentioned in previous articles [56–57] but must be tested in future research.

The present investigation suggests that pesticide regime should be taken in consideration when assessing the ecological impact of invasive species in a new environment. Studies of lethal and sublethal effects of compounds used in the environment involved, as well as effects on behaviour of invaders and intraguild interactions at a population level, should be envisaged. Our research also highlights the side effects of reduced-risk insecticides (novaluron) on beneficial organisms and potentially on invasive species and their ecological consequences. Novaluron is a wide spectrum insecticide highly toxic for the Asian ladybeetle [56–57], as well as for other natural enemies found orchards, such as the lacewing *Chrysoperla carnea* (Stephens) (Neur., Chrysopidae), the predatory plant bug *Deraeocoris brevis* (Uhler) (Hem., Miridae), the ladybeetle *Hippodamia convergens* Guérin-Méneville (Col., Coccinellidae), and the mite predators *Galendromus occidentalis* (Nesbitt) [71] and *Neoseiulus fallacis* (Garman) (Aca., Phytoseiidae) [72].

However, certain natural enemies are less susceptible to it, as it is the case of the twelve spotted ladybeetle [56–57] and the parasitoid *Aphelinus mali* (Hald.) (Hym., Aphelinidae) [71]. Accordingly, as mentioned in 4.1., impact of IGP on coccinellid assemblages, as well as on aphidophagous guilds facing pesticides will be determined by the variability of susceptibilities among guild members as well as neutralisation of toxic compounds by intraguild preys.

Moreover, insecticides can modify the composition of guilds, altering occurrences of competitors, which changes the dynamics of the guild and the frequency of encounters for IGP. Our results suggest that direct and indirect effects of insecticide treatments in agroecosystems are likely to have important impacts on ecosystem services of guilds, particularly on biocontrol by interfering with IGP.

The present study highlights the impact of an insecticide on an adaptive behaviour for a top predator, which is one of the key factors associated to the invasive status of the multicolored Asian ladybeetle worldwide. At an ecological level, our findings show that an insecticide might alter not only guild composition but also disturb intraguild interactions and consequently alter cascade effects in trophic systems.

## Acknowledgements

We thank Jill Vandermeerschen for her advice with statistics, Franz Vanoosthuyse for technical support, and Chloé Savoie, Maryse Pelletier, Mathieu Lemieux, and Marie Elen Dupuis for assisting with ladybeetle bioassays.

